# Critical role of type III interferon in controlling SARS-CoV-2 infection, replication and spread in primary human intestinal epithelial cells

**DOI:** 10.1101/2020.04.24.059667

**Authors:** Megan L. Stanifer, Carmon Kee, Mirko Cortese, Sergio Triana, Markus Mukenhirn, Hans-Georg Kraeusslich, Theodore Alexandrov, Ralf Bartenschlager, Steeve Boulant

**Author notes:** Corresponding authors Steeve Boulant, Ph.D., Department of Infectious Disease, Virology, Heidelberg University, Im Neuenheimer Feld 344, 69120 Heidelberg, Germany, Megan L. Stanifer, Ph.D., Department of Infectious Disease, Molecular Virology, Heidelberg University, Im Neuenheimer Feld 344, 69120 Heidelberg, Germany.

## Abstract

SARS-CoV-2 is an unprecedented worldwide health problem that requires concerted and global approaches to better understand the virus in order to develop novel therapeutic approaches to stop the COVID-19 pandemic and to better prepare against potential future emergence of novel pandemic viruses. Although SARS-CoV-2 primarily targets cells of the lung epithelium causing respiratory infection and pathologies, there is growing evidence that the intestinal epithelium is also infected. However, the importance of the enteric phase of SARS-CoV-2 for virus-induced pathologies, spreading and prognosis remains unknown. Here, using both colon-derived cell lines and primary non-transformed colon organoids, we engage in the first comprehensive analysis of SARS-CoV-2 lifecycle in human intestinal epithelial cells. Our results demonstrate that human intestinal epithelial cells fully support SARS-CoV-2 infection, replication and production of infectious *de-novo* virus particles. Importantly, we identified intestinal epithelial cells as the best culture model to propagate SARS-CoV-2. We found that viral infection elicited an extremely robust intrinsic immune response where, interestingly, type III interferon mediated response was significantly more efficient at controlling SARS-CoV-2 replication and spread compared to type I interferon. Taken together, our data demonstrate that human intestinal epithelial cells are a productive site of SARS-CoV-2 replication and suggest that the enteric phase of SARS-CoV-2 may participate in the pathologies observed in COVID-19 patients by contributing in increasing patient viremia and by fueling an exacerbated cytokine response.

## Introduction

*Coronaviridae* is a large family of single-stranded positive-sense enveloped RNA viruses that can infect most animal species (human as well as domestic and wild animals). They are known to have the largest viral RNA genome and are composed of four genera (Cui et al., 2019). Generally, infection by coronaviruses results in mild respiratory tract symptoms and they are known to be one of the leading causes of the common cold (Moriyama et al., 2020; Paules et al., 2020). However, in the last 18 years, we have witnessed the emergence of highly pathogenic human coronaviruses: the severe acute respiratory syndrome-related coronavirus (SARS-CoV-1), the Middle East respiratory syndrome-related coronavirus (MERS-CoV) and, at the end of 2019, the severe acute respiratory syndrome-related coronavirus-2 (SARS-CoV-2) (Lu et al., 2020). SARS-CoV-2 is responsible for the coronavirus-associated acute respiratory disease or coronavirus disease 19 (COVID-19) and represents a major global health threat and coordinated efforts are urgently needed to treat the viral infection and stop the pandemic.

Although SARS-CoV-2 primarily targets cells of the lung epithelium causing respiratory infection, there is growing evidence that the intestinal epithelium can also be infected. Multiple studies have reported gastro-intestinal symptoms such as diarrhea at the onset of the disease and have detected the prolonged shedding of large amounts of coronavirus genomes in the feces even after the virus was not detectable in oropharyngeal swabs (Wu et al., 2020b; Xiao et al., 2020; Xing et al., 2020; Xu et al., 2020b) (Wölfel et al., 2020). Although one study revealed the isolation of infectious virus particles from stool samples (Wang et al., 2020), to date, it remains unclear how many people shed infectious viruses in feces. Most critically, it remains unknown whether or not there is a possibility for fecal transmission of SARS-CoV-2 but multiple health agencies worldwide have highlighted this possibility. The presence of such a large amount of coronavirus genomes in feces is hardly explainable by a swallowing virus replicating in the throat or by a loss of barrier function of the intestinal epithelium which will allow the release of viruses or genomes from the inside of the body (circulation or *lamina propria*) to the lumen of the gut. Instead, it is likely due to an active replication in the intestinal epithelium. Recently, intestinal biopsies of SARS-CoV-2 infected patients clearly show the presence of replicating viruses in epithelial cells of the small and large intestine (Xiao et al., 2020). SARS-CoV-2 infection of the gastrointestinal tract is supported by the fact that ACE2, the virus receptor (Hoffmann et al., 2020), is expressed in intestinal epithelial cells (Zhao et al., 2020) (Lukassen et al., 2020) (Wu et al., 2020a) (Venkatakrishnan et al., 2020) and single cell sequencing analysis suggest that its expression is even higher on intestinal cells compared to lung cells (Xu et al., 2020a).This highlights that SARS-CoV-2 is not restricted to the lung but also infects the gastrointestinal tract. Importantly, many animal coronaviruses are well known to be enteric and are transmitted via the fecal-oral route (Wang et al., 2019)(Wang and Zhang, 2016). Additionally, the presence of human pathogenic coronaviruses in the gastrointestinal tract was previously reported for SARS-CoV-1 and MERS-CoV but remained seriously understudied (Leung et al., 2003; Wong et al.; Zhou et al., 2017). Although it is now clear that human coronaviruses, particularly SARS-CoV-2, are found in feces and can infect the gastrointestinal tract, the importance of its enteric phase for viremia, pathogenesis and patient prognosis remains unknown.

To combat the current pandemic of COVID-19 and to prepare for potential future emerging zoonotic coronaviruses, we need to gain a better understanding of the molecular basis of SARS-CoV-2 infection, replication and spread in a tissue-specific manner. Here, we engaged in studying SARS-CoV-2 infection of human intestinal cells. For this, we exploited both human intestinal epithelial cell lines and human organoid culture models to characterize how these cells support SARS-CoV-2 replication and spread and how they respond to viral infection. Direct comparison of both primary and transformed cells show that human intestinal epithelial cells fully support SARS-CoV-2 infection and *de-novo* production of infectious virus particles. Interestingly, viral infection elicited a robust intrinsic immune response where type III interferon mediated response was shown to be significantly more efficient at controlling SARS-CoV-2 replication and spread as compared to type I interferon. Importantly, human primary intestinal epithelial cells responded to SARS-CoV-2 infection by producing only type III interferon. Taken together, our data clearly highlight the importance of the enteric phase of SARS-CoV-2 and this should be taken into consideration when developing hygienic/containment measures, and antiviral strategies, and when determining patient prognosis.

## Results

### Efficient infection of human intestinal epithelial cells by SARS-CoV-2

As there is growing evidence that the gastro-intestinal tract is infected by SARS-CoV-2, we engaged in studying virus infection, replication and spread in human intestinal epithelial cells (IECs). First, SARS-CoV-2 (strain BavPat1) was propagated in the Green monkey cell line Vero (see Methods section). To detect viral infection, we used an antibody directed against a region of the nucleoprotein (NP) which is conserved between of SARS-CoV-1 and SARS-CoV-2. Additionally, we used the J2 antibody which detects double-stranded RNA (dsRNA) which is a hallmark of RNA virus replication (Targett-Adams et al., 2008). Cells positive for NP were found to be always positive for dsRNA; the NP signal was found to be dispersed within the cytosolic area whereas dsRNA were found in discrete foci likely corresponding to replication compartments (Harak and Lohmann, 2015)(Fig. S1A). Supernatants of infected Vero cells were collected at 48 hours post-infection (hpi) and the amount of infectious virus particles present was measured using a TCID50 approach on Vero cells (Fig. S1B). The colon carcinoma derived lines T84 and Caco-2 cells were then infected with SARS-CoV-2 at a MOI of 0.5 (as determined in Vero cells) and, at different time post-infection, cells were fixed and immunostained using the anti-NP and anti-dsRNA antibodies (Fig. 1A). Results show that SARS-CoV-2 infected Caco-2 cells were readily detected as early as 4hpi and, by 24 hpi, most of the cells were infected (Fig. 1B). Similar results were observed in the T84 cells, but detection of infection was slightly delayed compared to Caco-2 cells and, although the same amount of SARS-CoV-2 was used to infect these cells, only around 20% of the cells were found infected at 24 hpi (Fig. 1B). These observations were in agreement with the increase in viral genome copy numbers over time (Fig. 1C) and the release of infectious virus particles in the supernatant of infected T84 and Caco-2 cells (Fig. 1D). Interestingly, infection of IECs by SARS-CoV-2 was associated with the generation of an interferon (IFN)-mediated intrinsic immune response. Concomitant with the differences observed in virus replication and *de-novo* virus production observed between T84 and Caco-2 cells, T84 cells mounted a much stronger immune response compared to Caco-2 cells (Fig. 1E) although much less T84 cells were infected (Fig. 1B). All together these results show that IECs are readily infected by SARS-CoV-2 and that infection of Caco-2 cells lead to a weaker intrinsic immune response which is associated with more *de-novo* infectious virus production compared to T84 cells. This observation suggests that the IFN-mediated immune response controls SARS-CoV-2 infection in IECs.

**Figure 1.**
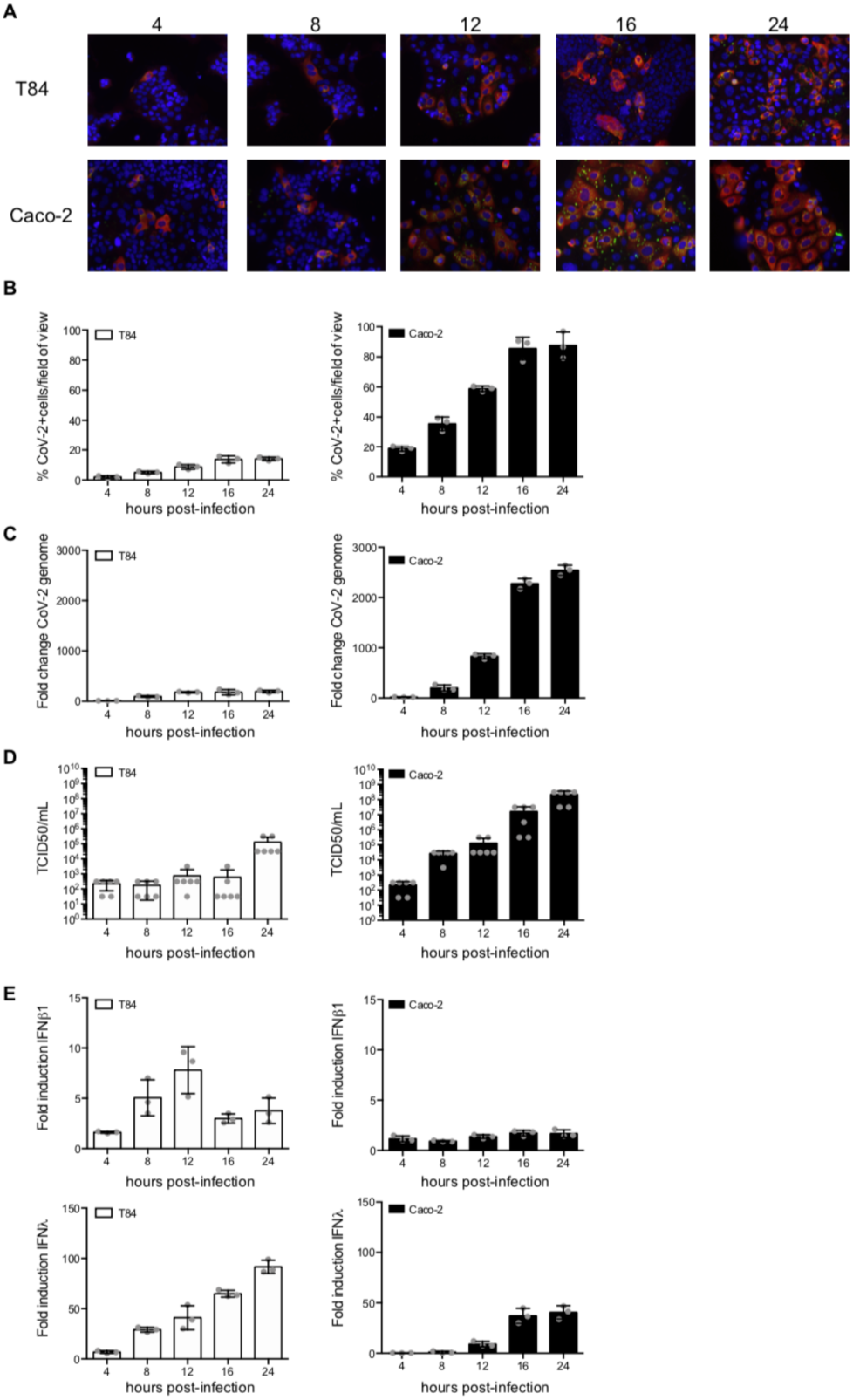
Human IECs support SARS-CoV-2 infection, replication and spread. Human colon carcinoma cells, T84 and Caco-2 cells, were infected with SARS-CoV-2 at a MOI of 0.5. (A) At indicated time points, cells were fixed and indirect immunofluorescence was performed against the viral NP protein (red) and dsRNA (green). Nuclei were stained with DAPI (blue). Experiments were performed in triplicate; representative images are shown. (B) Same as A except the number of SARS-CoV-2 positive cells was quantified in 10 fields of view for each time point. N=3. (C) At indicated time points, RNA was harvested and q-RT-PCR was used to evaluate the copy number of SARS-CoV-2 genome. N=3. (D) At indicated time points supernatants were collected from infected T84 and Caco-2 cells. The amount of *de-novo* viruses present in the supernatants was determined using a TCID50 assay on naïve Vero cells. N=3. (E) Same as C except the up regulation of IFNβ1 and IFNλ was evaluated. N=3. (B-) Error bars indicate standard deviation.

### Interferon hinders SARS-CoV-2 infection in human intestinal epithelial cells

To directly test the function of IFNs in controlling/restricting SARS-CoV-2 replication and spread in human IECS, T84 and Caco-2 cells were mock-treated or pre-treated with type I (IFNβ1) or type III (IFNλ) IFNs. 24 hrs post-treatment, cells were infected with SARS-CoV-2 at a MOI of 0.5 (as determined in Vero cells), in the presence or absence of IFNs (Fig. 2A). 24 hpi, cells were immunostained using both the anti-NP and anti-dsRNA antibodies. Results show that pre-treatment of cells with either IFNs significantly interfered with SARS-CoV-2 infection of both T84 and Caco-2 cells (Fig. 2B). This decrease of the number of SARS-CoV-2 infected cells pre-treated with either IFNs was associated with an inhibition of the increase in viral genome copy numbers (Fig. 2C) and with a significant decrease in the release of infectious *de-novo* virus particles (Fig. 2D). These results show that both IFNβ1 and IFNλ provided in *trans* can induce an efficient antiviral state in IECs preventing/controlling SARS-CoV-2 infection. This strongly suggests that the IFN-mediated immune response can control SARS-CoV-2 infection in human IECs.

**Figure 2.**
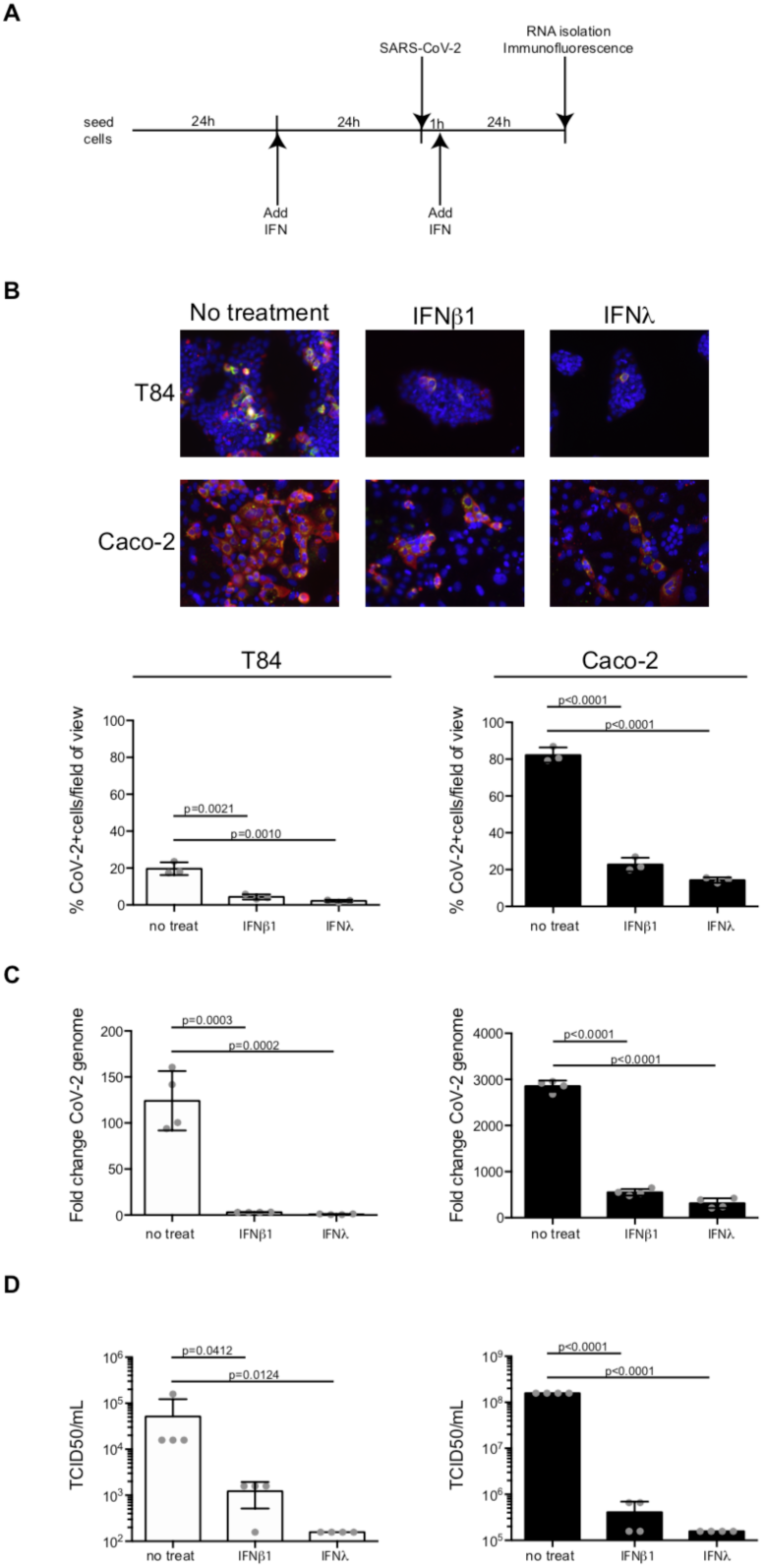
IFNβ and IFNλ control SARS-CoV-2 infection of human IECs. (A) Schematic describing the experimental set-up. (B) T84 and Caco-2 cells were pre-treated with IFNs 24 hrs prior to infection. SARS-CoV-2 was added to cells for one hour. Upon removal of the virus, new media containing IFNs was added to cells. 24 hpi virus infection was evaluated by indirect immunofluorescence for the viral NP protein (red) and dsRNA (green). Nuclei were stained with DAPI (blue). Experiments were performed in triplicate; representative images are shown. The number of SARS-CoV-2 positive cells was quantified in ten fields of view for each time point. (C) Same as B except RNA was harvested and q-RT-PCR was used to evaluate the copy number of SARS-CoV-2 genome. N=3. (D) 24 hpi supernatants were collected from infected T84 and Caco-2 cells. The amount of *de-novo* viruses present in the supernatants was determined using a TCID50 assay on naïve Vero cells. N=3 (B-D) Error bars indicate standard deviation. P values were determined by unpaired t-test.

### Interferon-mediated intrinsic immune response controls SARS-CoV-2 infection in human intestinal epithelial cells

To address whether the endogenous levels of IFNs generated by IECs controls SARS-CoV-2 replication and spread, we exploited our previously reported IECs depleted of either the type I IFN receptor (IFNAR1) (AR-/-), the type III IFN receptor (IFNLR1) (LR -/-) or depleted of both IFN receptors (dKO). To control that our cells were functionally knocked-out for the type I IFN (AR-/-) and/or the type III IFN receptor (LR-/-), T84 cells were treated with either IFN and the production of the IFN stimulated gene IFIT1 was evaluated. As expected, type I IFN receptor knock-out cells (AR-/-) only responded to IFNλ, whereas type III IFN receptor knock-out cells (LR-/-) only responded to IFNβ1 (Fig. S2). The IFN receptor double knock-out cells (dKO) did not respond to either IFN (Fig. S2). WT or IFN receptor knock-out T84 cells were infected with SARS-CoV-2 at a MOI of 0.1 (as determined in T84 cells). 24 hpi, cells were immunostained using the anti-NP antibody and the number of infected cells was quantified using fluorescent microscopy (Fig. 3A). Results showed that depletion of the type I IFN receptor (AR-/-) resulted in a slight increase in the number of infected cells. Importantly, depletion of the type III IFN receptor (LR-/-) resulted in a massive increase of cell infectivity by a factor of around seven. Similar results were obtained when both the type I and the type III IFN receptors were depleted (dKO) (Fig. 3B). Interestingly, this increase in the number of infected cells upon depletion of the type III IFN receptor (LR-/-) was associated with a significant increase in viral genome copy numbers (Fig. 3C) and with a three orders of magnitude increase in *de-novo* infectious virus production (Fig. 3D). Together these results suggest that the type III IFN-mediated immune response actively participates in controlling SARS-CoV-2 infection in human IECs.

**Figure 3.**
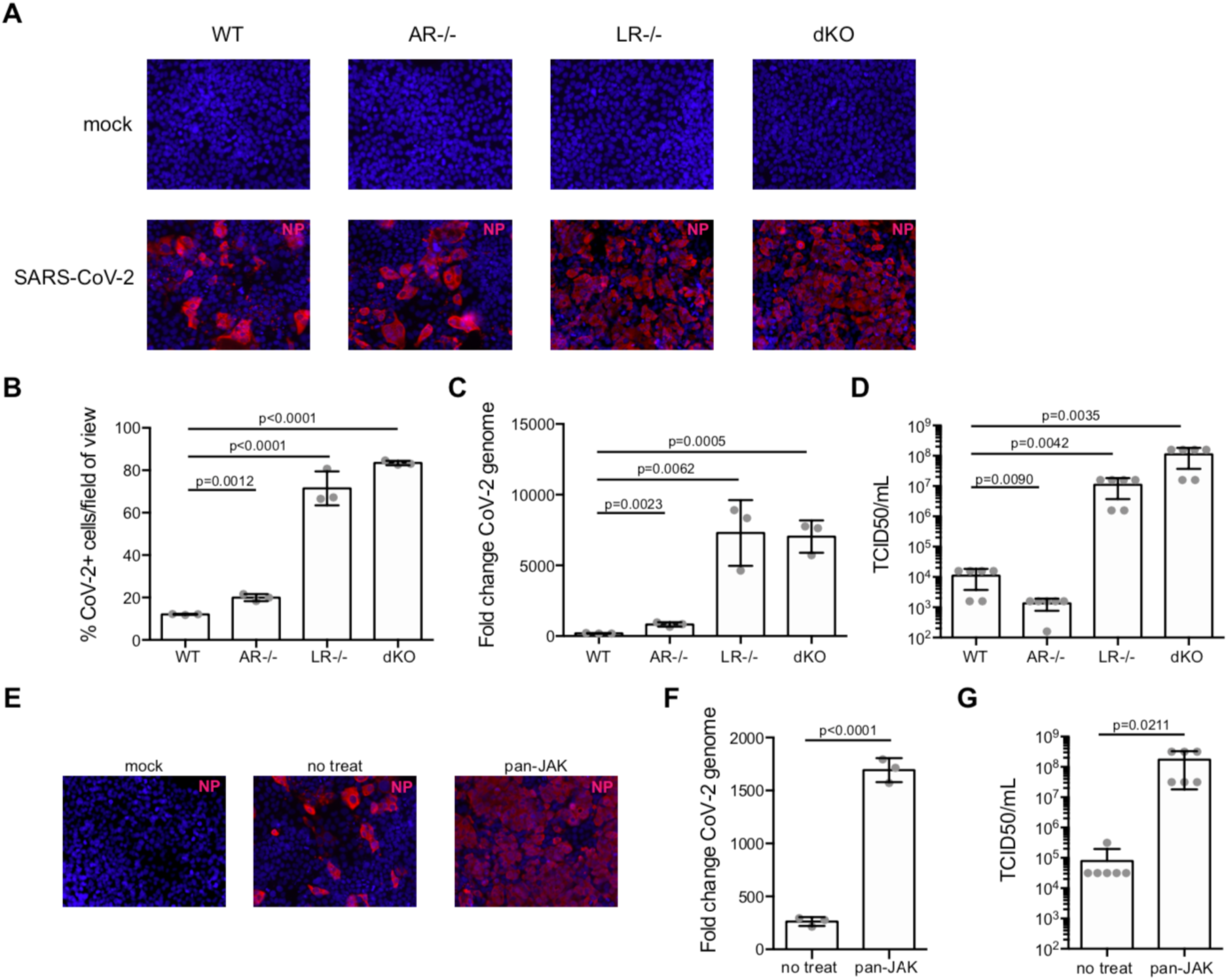
Type III IFN receptor controls SARS-CoV-2 replication in human IECs. (A-D) Wild type T84 cells and T84 cells depleted of the type I IFN receptor (AR-/-), the type III IFN receptor (LR-/-) or both (dKO) were infected with SARS-CoV-2 at an MOI of 0,1. (A) 24 hpi, infection was analyzed by indirect immunofluorescence of the viral NP protein (red). Nuclei were stained with DAPI (blue). Experiments were performed in triplicate; representative images are shown. (B) Same as A except the number of SARS-CoV-2 positive cells was quantified in 10 fields of view for each cell type. N=3. (C) 24 hpi, RNA was harvested and the change in the copy number of SARS-CoV-2 genome was evaluated by q-RT-PCR. N=3. (D). 24 hpi, supernatants were collected from all cell types. The amount of *de-novo* viruses present in the supernatants was determined using a TCID50 assay on naïve Vero cells. N=3. (E-G) Wild type T84 cells were mock treated or pre-treated with pyridine-6, a pan-JAK inhibitor, for 2 hours prior to SARS-CoV-2 infection. Following infection, the pan-JAK inhibitor was replaced and maintained for the course of the infection. (E) 24 hpi, infection was analyzed by indirect immunofluorescence of the viral NP protein (red). Nuclei were stained with DAPI (blue). Experiments were performed in triplicate; representative images are shown. (F) 24 hpi, RNA was harvested and the change in the copy number of SARS-CoV-2 genome was evaluated by q-RT-PCR. N=3. (G) 24 hpi, supernatants were collected from mock-treated or pan-JAK treated cells. The amount of *de-novo* viruses present in the supernatants was determined using a TCID50 assay on naïve Vero cells. N=3. (B-D; F-G) Error bars indicate standard deviation. P values were determined by unpaired t-test.

To unambiguously address the importance of the IFN-mediated antiviral response, we used the pan-JAK inhibitor (pyridone-6) to inhibit the STAT1 phosphorylation activation and block the production of interferon stimulated genes (ISGs). As we previously reported, treatment of T84 cells with the pan-JAK inhibitor fully inhibits signal transduction downstream both the type I and type III IFN receptors ((Pervolaraki et al., 2017) and data not shown). Mock and pyridone-6 pre-treated T84 cells were infected with SARS-CoV-2 for 24hrs and analyzed using fluorescence microscopy following immunostaining using the anti-NP antibody. Results show both a significant increase in the number of infected cells (Fig. 3E) and an increase of viral genome copy number in cells treated with the pan-JAK inhibitor (Fig.3F). Importantly, and in agreement with the results observed in cells depleted of the type III IFN receptor, this increase in infectivity was also associated with an increase in infectious *de-novo* virus particle production (Fig. 3G). All together, these results strongly support a model where the type III IFN mediated signaling controls SARS-CoV-2 infection in human intestinal epithelial cells.

### Human primary intestinal organoids support SARS-CoV-2 infection, replication and spread

To address whether primary human IECs can be infected by SARS-CoV-2 and support *de-novo* virus production, we used human colon derived organoids from two distinct donors. Intact ultrastructural organization and differentiation to all cell types (e.g. enterocytes, Goblet cells, enteroendocrine cells, and stem cells) was confirmed using confocal fluorescent microscopy (Fig. 4A) and quantitative-RT-PCR against cell type specific transcripts (Fig. 4B). As quantification of the number of infected cells in 3D organoids is very challenging, we exploited previously established protocols to differentiate and infect human intestinal organoids in two dimensions with viruses (Ettayebi et al., 2016; Stanifer et al., 2020). Non-differentiated organoids were seeded on human collagen-coated iBIDI chambers. At 24 hrs post-seeding, differentiation was induced by removal of Wnt3a and reducing the amounts of R-spondin and Noggin for four days. Upon full differentiation, organoids were infected with SARS-CoV-2. At 24 hpi, the infection was analyzed by immunostaining using the anti-NP and anti-dsRNA antibodies and by quantitative RT-PCR (Fig. 4C). Results show that independent of the donor, colon organoids were readily infected by SARS-CoV-2 as noted by the presence of cells positive for both NP and dsRNA (Fig. 4D). Quantification revealed that around 8-13% of cells were infected in each donor (Fig. 4E). This infection was associated with an increase of viral genome copy number (Fig. 4F). Interestingly, infection of organoids led to no type I interferon (IFNβ1) production but an extremely large up regulation of type III IFN (IFNλ) (Fig. 4G and Fig. S3). To determine if exogenously added IFNs could prevent SARS-CoV-2 infection, colon organoids were mock or pre-treated with IFNβ1 and IFNλ and then subsequently infected with SARS-CoV-2 for 24 hrs. Results show that pre-treatment of colon organoids with both IFNβ1 and IFNλ significantly impaired infection (Fig. 4H). This was associated with a reduction of SARS-CoV-2 genome copy numbers (Fig. 4I) and a decrease in infectious *de-novo* virus particle production (Fig. 4J). All together these results show that human colon organoids can support SARS-CoV-2 infection, replication and spread and that the type III IFN response plays a critical role in controlling virus replication.

**Figure 4.**
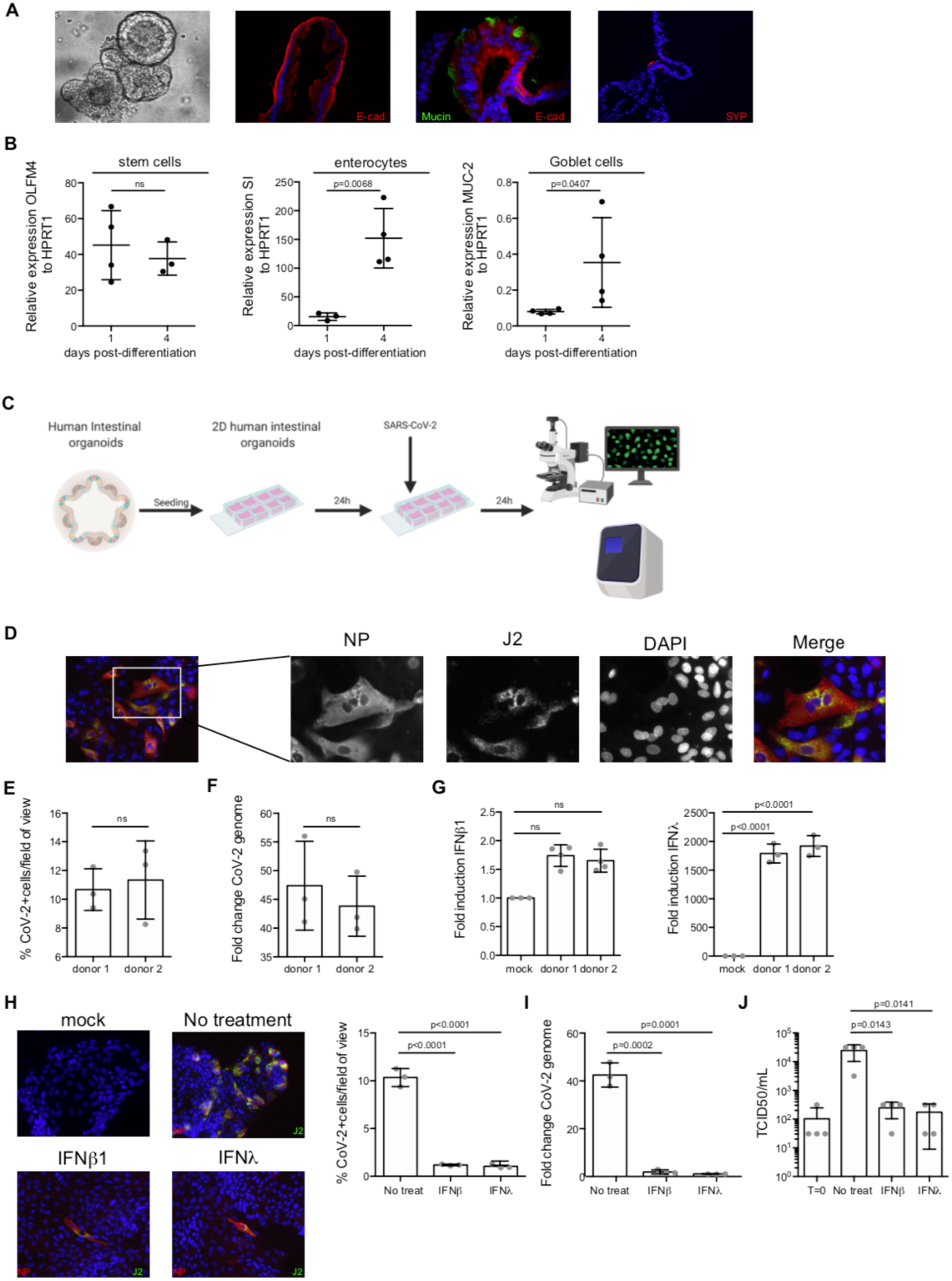
Human colon organoids support SARS-CoV-2 infection, replication and spread. (A-B) Validation of proper differentiation of human colon organoids. (A) Bright field and immunofluorescence images showing human colon organoids that express markers for enterocytes (E-cad), Goblet cells (mucin) and enteroendocrine cells (SYP). Representative images are shown. (B) Colon organoids were differentiated prior to infection with SARS-CoV-2. Differentiation was confirmed by q-RT-PCR for stem cell markers (OLFM4), enterocytes (SI), Goblet cells (MUC-2). N=3. (C) Schematic describing method for infection of 2D colon organoids with SARS-CoV-2. (D-G) Colon organoids were seeded in 2D and differentiated prior SARS-CoV-2 infection. 24 hpi, cells were analyzed for virus replication and immune response. (D) Infection was analyzed by indirect immunofluorescence of the viral NP protein (red) and dsRNA (green). Nuclei were stained with DAPI (blue). Experiments were performed in triplicate; representative images are shown. (E) The number of SARS-CoV-2 positive cells was quantified in 10 fields for each donor. N=3. (F) RNA was harvested and the change in the copy number of SARS-CoV-2 genome was evaluated by q-RT-PCR. N=3. (G) same as F, except the up regulation of IFNβ1 and IFNλ was evaluated. N=3. (H-J) Colon organoids were pre-treated with IFNs 24 hrs prior to infection. SARS-CoV-2 was added to organoids for one hour. Upon removal of the virus, new media containing IFNs was added to cells. (H) 24 hpi, virus infection was evaluated by indirect immunofluorescence for the viral NP protein (red) and dsRNA (green). Nuclei were stained with DAPI (blue). Experiments were performed in triplicate; representative images are shown. The number of SARS-CoV-2 positive cells was quantified in ten fields of view for each time point. (I) RNA was harvested and q-RT-PCR was used to evaluate the copy number of SARS-CoV-2 genome. N=3. (J) supernatants were collected from infected organoids. The amount of *de-novo* viruses present in the supernatants was determined using a TCID50 assay on naïve Vero cells. N=3. Error bars indicate standard deviation. P values were determined by unpaired t-test.

## Discussion

Increasing numbers of clinical reports detail that COVID-19 patients not only have severe respiratory symptoms but also show indications of gastrointestinal pathologies. To efficiently combat the pandemic, it is necessary to fully appreciate the complexity of the disease and this requires that the virus life cycle and its interaction with the host are characterized for each site of infection (*e*.*g*. lung *vs*. gut). Here, we report that human intestinal epithelial cells, including primary non-transformed epithelial cells, fully support SARS-CoV-2 infection, replication and spread. Interestingly, infection of human primary intestinal epithelial cells by SARS-CoV-2 induces a robust intrinsic immune response characterized by the production of type III IFNs but not type I IFN. Pre-treatment of intestinal epithelial cells with exogenous IFNs prior to SARS-CoV-2 infection results in a major drop of the number of infected cells, a reduction of viral replication and a significant decrease in the production of infectious *de-novo* virus particles. Concomitantly, inhibition of type III IFN signaling by genetic ablation of its specific receptor results in a significant increase of SARS-CoV-2 replication and secretion. Together our results demonstrate that human intestinal epithelial cells are a productive site of SARS-CoV-2 replication and this may have critical implications for the etiology of both SARS-CoV-2 infection and for the pathologies observed in COVID-19 patients. Additionally, our work highlights the central role of type III IFN in controlling SARS-CoV-2 at the intestinal epithelium.

Human primary intestinal epithelial cells support robust replication of SARS-CoV-2 and secretion of infectious *de-novo* virus particles. We found that around 10% of the cells are infected in a human colon derived organoid leading to a modest but significant increase in viral genome copy number (Fig. 4F) and *de-novo* infection virus particles (Fig. 4J) over time. This modest replication of SARS-CoV-2 is explained by the fact that (i) only a small fraction of the cells is infected by SARS-CoV-2, and (ii) as shown in this work, IFNs are potent inhibitors of SARS-CoV-2 replication (Fig. 2 and 3) and organoids are highly immunoresponsive upon viral infection (Stanifer et al., 2020) (Fig. 4G). It is well known that *in vivo* intestinal epithelium cells are less immunoresponsive because of the gut microenvironment (*e*.*g*. microbiota and tissue specific immune cells) and as such they will very likely show a severely dampened immune response allowing for an even greater SARS-CoV-2 replication. Importantly, the intestinal epithelium is the largest organ in the body and even if only a few percent of the cells are infected it will result in the generation of an extremely large amount of *de-novo* viruses.

Analysis of the single-cell RNA sequencing data from the Colon Atlas revealed that only 3.8% of human colon epithelial cells express very low levels of the SARS-CoV-2 receptor ACE-2 (Fig. S4A-E). This is very different to the small intestine where ACE-2 appears to be more expressed (Qi et al., 2020). This low ACE-2 level could explain why we have a small percentage of infected cells in our colon organoids (Fig. 4E). On the contrary, TMPRSS2 seems to be not a limiting factor in the colon (Fig. S4C-E). Interestingly, q-RT-PCR and western blot analysis do not support single cell analysis and clearly show that ACE-2 is expressed in both our carcinoma derived lines and our colon organoids (Fig. S4F-G). The discrepancy between the single-cell RNA sequencing and classical molecular and biochemical approaches is likely the results of (i) the sequencing being not deep enough to detect ACE-2 in individual cells and (ii) RNA expression not necessarily matching the protein expression levels. These observations highlight that although analyzing data from single-cell RNA sequencing atlases could be very informative, their findings should be validated in tissues as they may be mis-leading or miss important sites of virus replication.

The human colon carcinoma Caco-2 cells produce very large amounts of infectious viral particles (between 108 and 109 infectious particles per mL). This is higher than the titers obtained in Vero cells which are commonly used to isolate and propagate SARS-CoV-2 (Harcourt et al., 2020) and where we routinely obtained titers of 105 and 106 infectious particles per mL in Vero cells (Fig. S1B). Interestingly, in a study comparing 13 different human cell lines, Caco-2 cells were the only cell type found to support SARS-CoV-1 replication and, compared to green monkey cells, Caco-2 cells were very efficient in producing infectious *de-novo* virus particles and did not show cytopathic effects (Mossel et al., 2005). These observations that Caco-2 are excellent culture models supporting both SARS-CoV-2 and SARS-CoV-1 further highlight the potentially central role of intestinal epithelial cells in COVID-19 patients. Intriguingly, T84 cells, which are also colon-carcinoma derived cells supported SARS-CoV-2 infection, replication and spread but to a much lesser extent compared to Caco-2 cells. Analysis of the intrinsic immune response generated upon infection of both cell lines revealed that T84 cells are more immunoresponsive compared to Caco-2 cells. In light of the results obtained by pretreating cells with exogenous IFNs, we propose that T84 cells, by being able to mount a stronger and faster immune response compared to Caco-2 cells can better restrict SARS-CoV-2 infection. This model is fully supported by our IFN receptor knock-out T84 cells, in which virus replication and *de-novo* virus production are drastically increased to levels similar to the ones observed in Caco-2 cells.

Exogenously added IFNs (both type I and type III IFNs) induce an antiviral state in our human intestinal epithelial cells, thereby restricting SARS-CoV-2 replication in these cells. Similar observations were made with type I IFN in Vero cells (Mantlo et al., 2020). Interestingly, infection of the Calu-3 human lung epithelial cells by SARS-CoV-2 seem to also mount an immune response (Lokugamage et al., 2020), whereas, in A549 cells, in ferret models and in human lung tissues, SARS-CoV-2 infection lead to the induction of a restricted and muted immune response lacking the induction of type I and type III IFN (Blanco-Melo et al., 2020) (Chu et al.). The reason for the difference in the amount of IFN made upon infection of lung *vs*. intestinal epithelial cells is currently unclear. Like intestinal epithelial cells (Pervolaraki et al., 2017, 2018; Pott et al., 2011), lung epithelial cells are normally highly immunoresponsive and make IFNs upon viral infection (Crotta et al., 2013; Ye et al., 2019). The lack of IFN in lung epithelial cells following SARS-CoV-2 infection might be specific for this virus in this tissue (Blanco-Melo et al., 2020) as SARS-CoV-1 induces IFN production in infected lung tissue (Chu et al.). These differences should be properly addressed in future studies. Importantly, our work shows that, compared to the airway epithelium, the intestinal epithelium produces a typical antiviral response. This highlights that host/pathogen interaction should be considered in a tissue specific manner as different cellular responses and viral countermeasures might be established between the lung, the gut and other organs.

Interestingly, in human colon organoids we observed that only type III IFN is made upon SARS-CoV-2 infection, although human intestinal organoids are capable of making both type I and III IFN upon enteric virus infection (Pervolaraki et al., 2018; Stanifer et al., 2020). The lack of type I induction appears to be specific to SARS-CoV-2 and it is likely that this virus encodes a specific antagonist which counteracts the production of type I IFN only. However, further studies are necessary to prove this novel concept.

We propose that the gut is an active site of replication for SARS-CoV-2 and this could account for the viremia observed in COVID-19 patients and for the presence of large amounts of SARS-CoV-2 genomes in the feces. The origin of the replicating SARS-CoV-2 in the intestinal epithelium is still not clear. To date only one paper reported the isolation of infectious virus from stool samples (Wang et al., 2020). Further characterization of the SARS-CoV-2 enteric lifecycle is necessary to determine whether the viral infection observed in the gut is due to fecal/oral transmission or is a manifestation of virus spreading from the lung to the gut. In the context of the gut we foresee that at the onset of SARS-CoV-2 infection, human intestinal epithelial cells will mount an antiviral response through the type III IFN signaling pathway. As immune cells participate in mounting an innate immune response, type I IFN will be secreted from these cells and will be able to act on intestinal epithelial cells further reinforcing their antiviral state against SARS-CoV-2. In respect to the severe pathologies observed in the lung, which are believed to be caused by a cytokine storm, the findings that lung epithelial cells mount a muted immune response upon SARS-CoV-2 infection suggests that the cytokines are coming from an alternative source. This source could be from local immune cells but also from the gastro-intestinal mucosa given the large immune response generated by primary human intestinal epithelial cells. As a matter of fact, many cytokines are released from epithelial cells towards the *lamina propria* (tissue side) (Stanifer et al., 2016), and will quickly enter the circulation to potentially fuel and promote the inflammation and pathology observed in the lung.

In conclusion, the gastro-intestinal tract is an active site of SARS-CoV-2 replication and this should be considered when developing antiviral strategies as it may participate in the viremia and potentially reinfection. From a patient prognosis point of view, the severity of the disease should be correlated with the extent of the enteric replication.

## STAR⋆Methods

### Cell and viruses

T84 human colon carcinoma cells (ATCC CCL-248) were maintained in a 50:50 mixture of Dulbecco’s modified Eagle’s medium (DMEM) and F12 (GibCo) supplemented with 10% fetal bovine serum and 1% penicillin/streptomycin (Gibco). Caco-2 human colorectal adenocarcinoma (ATCC HTB-37) and Vero E6 (ATCC CRL 1586) were maintained in Dulbecco’s modified Eagle’s medium (DMEM) (GibCo) supplemented with 10% fetal bovine serum and 1% penicillin/streptomycin (Gibco). SARS-CoV-2 (strain BavPat1) was obtained from Prof. Christian Drosten at the Charité in Berlin, Germany and provided via the European Virology Archive. The virus was amplified in Vero E6 cells.

### Human organoid cultures

Human tissue was received from colon resection from the University Hospital Heidelberg. This study was carried out in accordance with the recommendations of the University Hospital Heidelberg with informed written consent from all subjects in accordance with the Declaration of Helsinki. All samples were received and maintained in an anonymized manner. The protocol was approved by the “Ethics commission of the University Hospital Heidelberg” under the protocol S-443/2017. Stem cells containing crypts were isolated following 2 mM EDTA dissociation of tissue sample for 1 h at 4°C. Crypts were spun and washed in ice cold PBS. Fractions enriched in crypts were filtered with 70 μM filters and the fractions were observed under a light microscope. Fractions containing the highest number of crypts were pooled and spun again. The supernatant was removed, and crypts were re-suspended in Matrigel. Crypts were passaged and maintained in basal and differentiation culture media (see table 1).

**Table 1.**

| *Compound* | *Final concentration* |
| --- | --- |
| **Basal media** |  |
| Ad DMEM/F12<br>+GlutaMAX<br>+HEPES<br>+P/S |  |
| Wnt3A | 50% by volume |
| B27 | 1:50 |
| N2 | 1:100 |
| N-acetyl-cysteine | 1mM |
| R-spondin | 10% by volume |
| Noggin | 100ng/mL |
| EGF | 50ng/mL |
| Gastrin | 10mM |
| Nicotinamide | 10mM |
| A83-01 | 500nM |
| Sb202190 | 10uM |
| **Differentiation Media** |  |
| Ad DMEM/F12<br>+GlutaMAX<br>+HEPES<br>+P/S |  |
| B27 | 1:50 |
| N2 | 1:100 |
| N-acetyl-cysteine | 1mM |
| R-spondin | 5% by volume |
| Noggin | 50ng/mL |
| EGF | 50ng/mL |
| Gastrin | 10mM |
| A83-01 | 500nM |
| Sb202190 | 10uM |

### Antibodies and Inhibitors

Mouse monoclonal antibody against SARS-CoV NP (Sino biologicals MM05), mouse monoclonal against J2 (scions), mouse monoclonal against E-cadherin (BD Transductions #610181), and rabbit polyclonal anti-Mucin-2 (Santa Graz Biotechnology# sc-15334) were used at 1:500 for immunofluorescence. Rabbit polyclonal ACE-2 antibody (Abcam) was used at 1:500 and mouse polyclonal actin was used at 1:5000 for western blot. Secondary antibodies were conjugated with AF488 (Molecular Probes), AF568 (Molecular Probes), CW800 (Li-Cor) or CW800 (Li-Cor) directed against the animal source. Human recombinant IFN-beta1a (IFNβ) was obtained from Biomol (#86421) was used at a final concentration of 2,000IU/mL. Recombinant human IFNλ 1 (IL-29) (#300-02L), IFNλ 2 (IL28A) (#300-2K) and IFNλ 3 (IL-28B) (#300-2K) were purchased from Peprotech and were used at a concentration of 100ng/mL each to make a final concentration of 300ng/mL, Pyridone 6 (Calbiochem #420099-500)was used at a final concentration of 2uM.

### Viral infections

All SARS-CoV-2 infections were performed the MOI indicated in the text. Media was removed from cells and virus was added to cells for 1 hour at 37°C. Virus was removed, cells were wash 1x with PBS and media or media containing inhibitors/cytokines was added back to the cells.

### 2D organoid seeding

8-well iBIDI glass bottom chambers were coated with 2.5% human collagen in water for 1 h prior to organoids seeding. Organoids were collected at a ratio of 100 organoids/transwell. Collected organoids were spun at 450g for 5 mins and the supernatant was removed. Organoids were washed 1X with cold PBS and spun at 450g for 5 mins. PBS was removed and organoids were digested with 0.5% Trypsin-EDTA (Life technologies) for 5 mins at 37°C. Digestion was stopped by addition of serum containing medium. Organoids were spun at 450g for 5 mins and the supernatant was removed and organoids were re-suspended in normal growth media at a ratio of 250 μL media/well. The collagen mixture was removed from the iBIDI chambers and 250 μL of organoids were added to each well.

### RNA isolation, cDNA, and Qpcr

RNA was harvested from cells using RNAeasy RNA extraction kit (Qiagen) as per manufactures instructions. cDNA was made using iSCRIPT reverse transcriptase (BioRad) from 250 ng of total RNA as per manufactures instructions. q-RT-PCR was performed using iTaq SYBR green (BioRad) as per manufacturer’s instructions, TBP or HPRT1 were used as normalizing genes. Primer used:

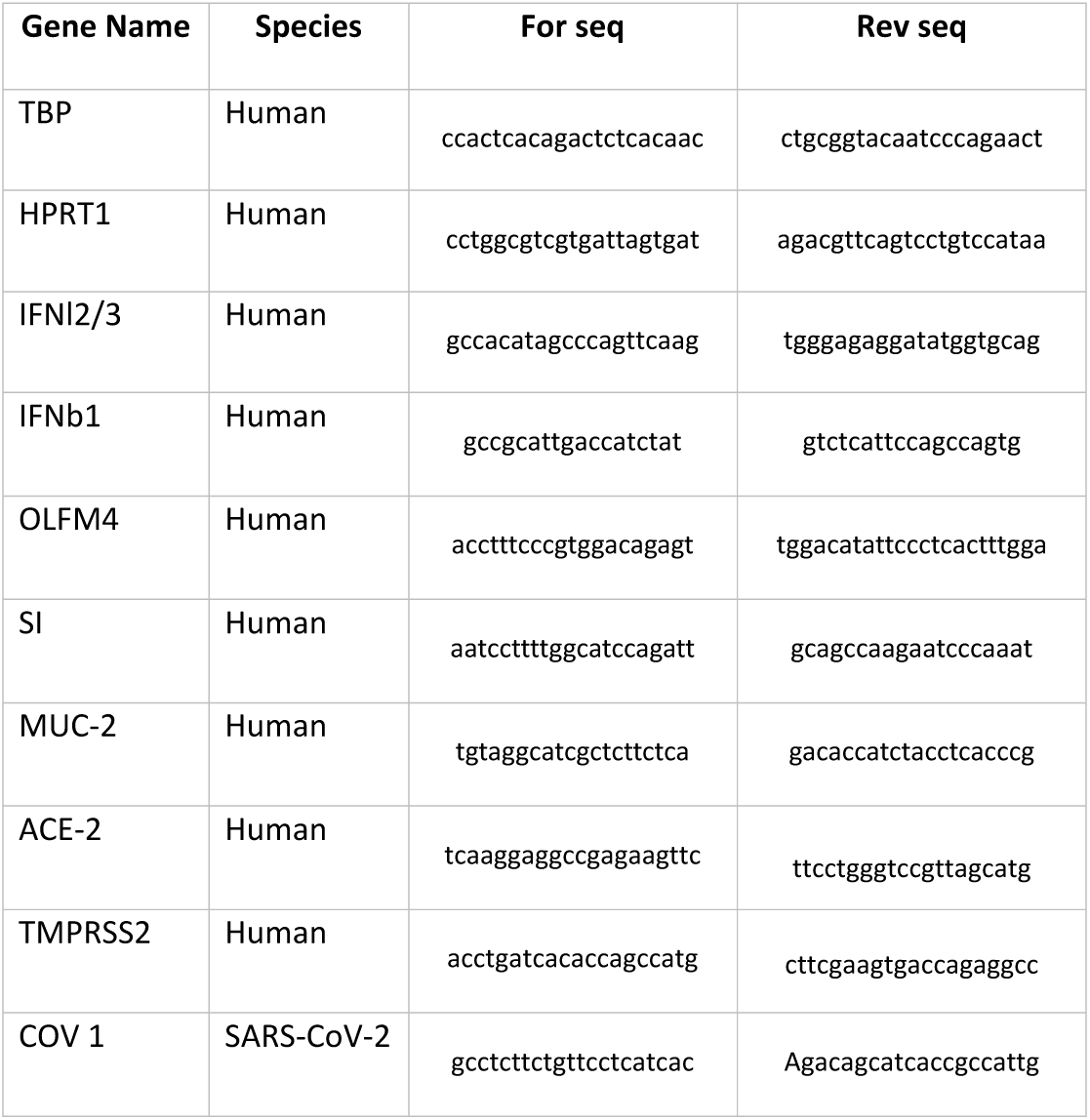

### Indirect Immunofluorescence Assay

Cells seeded on iBIDI glass bottom 8-well chamber slides. At indicated times post-infection, cells were fixed in 4% paraformaldehyde (PFA) for 20 mins at room temperature (RT). Cells were washed and permeabilized in 0.5% Triton-X for 15 mins at RT. Primary antibodies were diluted in phosphate-buffered saline (PBS) and incubated for 1h at RT. Cells were washed in 1X PBS three times and incubated with secondary antibodies and DAPI for 45 mins at RT. Cells were washed in 1X PBS three times and maintained in PBS. Cells were imaged by epifluorescence on a Nikon Eclipse Ti-S (Nikon).

### In-cell Western

20,000 Vero E6 cells were seeded per well into a 96-well dish 24h prior to infection. 100uL of harvested supernatant was added to the first well. Seven 1:10 dilutions were made (all samples were performed in triplicate). Infections were allowed to proceed for 24h. 24h post-infection cells were fixed in 2% PFA for 20mins at RT. PFA was removed and cells were washed twice in 1X PBS and then permeabilized for 10mins at RT in 0.5% Triton-X. Cells were blocked in a 1:2 dilution of Licor blocking buffer (Licor) for 30mins at RT. Cells were stained with 1/1000 dilution anti-dsRNA (J2) for 1h at RT. Cells were washed three times with 0.1% Tween in PBS. Secondary antibody (anti-mouse CW800) and DNA dye Draq5 (Abcam) were diluted 1/10,000 in blocking buffer and incubated for 1h at RT. Cells were washed three times with 0.1% Tween/PBS. Cells were imaged in 1X PBS on a LICOR (Li-Cor) imager.

### Western blot

At time of harvest, media was removed, cells were rinsed once with 1X PBS and lysed with 1X RIPA (150 mM sodium chloride, 1.0% Triton X-100, 0.5% sodium deoxycholate, 0.1% sodium dodecyl sulphate (SDS), 50 mM Tris, pH 8.0 with phosphatase and protease inhibitors (Sigma-Aldrich)) for 5 mins at room temperature (RT). Lysates were collected and equal protein amounts were separated by SDS-PAGE and blotted onto a nitrocellulose membrane by wet-blotting (Bio-Rad). Membranes were blocked with LiCor blocking buffer (LiCor) for one hour at RT. Primary antibodies were diluted in blocking buffer and incubated overnight at 4°C. Membranes were washed 3X in TBS-T for 5 mins at RT. Secondary antibodies were diluted in blocking buffer and incubated at RT for 1 hour with rocking. Membranes were washed 3X in TBS-T for 5 mins at RT and scanned on a LiCor scanner.

### Re-processing of the single-cell RNAseq Human Colon Atlas

The data from the single-cell transcriptome human colon atlas (Smillie et al., 2019) was obtained from the broad institute Single Cell Portal accession number SCP259 (https://singlecell.broadinstitute.org/single_cell/study/SCP259). Expression matrices of the epithelial subset of 30 samples were imported and individually analyzed using the Seurat software, version 3.1.4 (https://github.com/satijalab/seurat). Quality filtering was performed, and cells having fewer than 500 genes and more than 30% of UMI count mapped to mitochondrial genes were discarded. Consecutively, the resulting datasets were normalized, scaled and high-variance genes genes were selected. Reciprocal PCA-based data integration was done to merge the samples. Afterwards, the resulting batch-corrected counts were used for calculating PCA-based dimensionality reduction and unsupervised Louvain clustering. Furthermore, UMAP visualization was calculated using 50 neighboring points for the local approximation of the manifold structure. Cell type annotation was based on the unsupervised clustering and the metadata provided by the colon atlas (Smillie et al., 2019).

### Statistics

Unless otherwise stated, statistical analysis was performed by a two-tailed unpaired t test using the GraphPad Prism software package. All samples were analyzed without blinding or exclusion of samples.

## Acknowledgments

This work was supported by a research grant from Chica and Heinz Schaller Foundation and Deutsche Forschungsgemeinschaft (DFG): Project number 240245660 (Project 14 of SFB 1129), project number 278001972 (Project A09 of TRR186), project number 415089553 (Heisenberg grant) and project number 272983813 (project 22 of TRR179) to SB. DFG project number 416072091 to MS. ST was supported by the Darwin Trust of Edinburgh. Work of R.B. was supported by the Deutsche Forschungsgemeinschaft (DFG) – Projektnummer 272983813 – TRR 179, Projektnummer 240245660 – SFB 1129 and BA1505/8-1. TA is supported by the European Research Council (grant agreement773089). We would like to acknowledge Vibor Laketa and the Infectious Diseases Imaging Platform (IDIP) at the Center for Integrative Infectious Disease Research, Heidelberg, Germany, for support with image acquisition and analysis. We also thank Christian Drosten at the Charité, Berlin and the European Virus Archive (EVAg) for the provision of the SARS-CoV-2 strain BavPat1.

## Supplementary Figure legends

**Figure S1:**
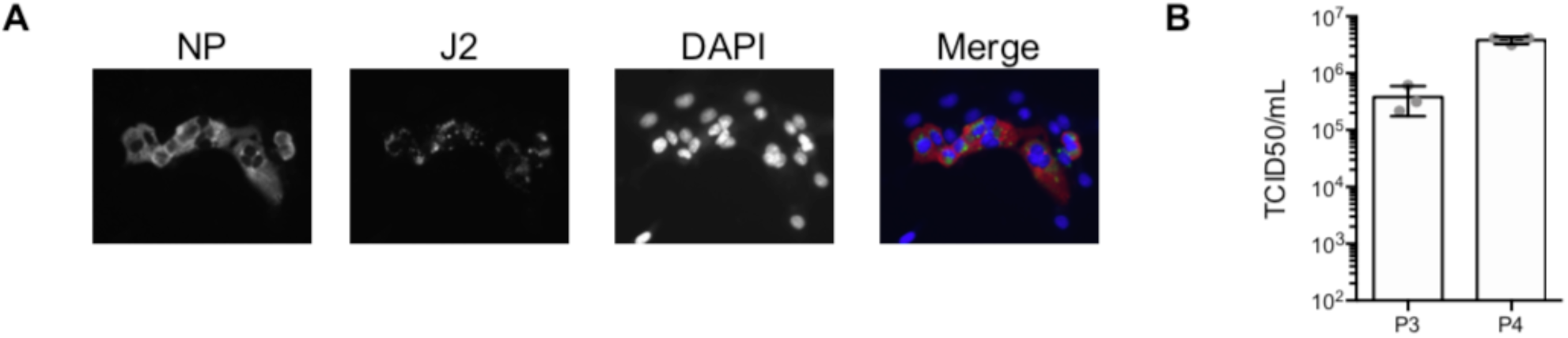
Vero cells support SARS-CoV-2 infection. (A) Vero cells were infected with SARS-CoV-2 and 24 hpi viral replication was evaluated by indirect immunofluorescence for the viral NP protein (red) and dsRNA (green). Nuclei were stained with DAPI (blue). Experiments were performed in triplicate, representative images are shown. (B) Viral titers from the passage P3 and P4 stocks was determined by TCID50. N=3.Error bars indicate standard deviation.

**Figure S2.**
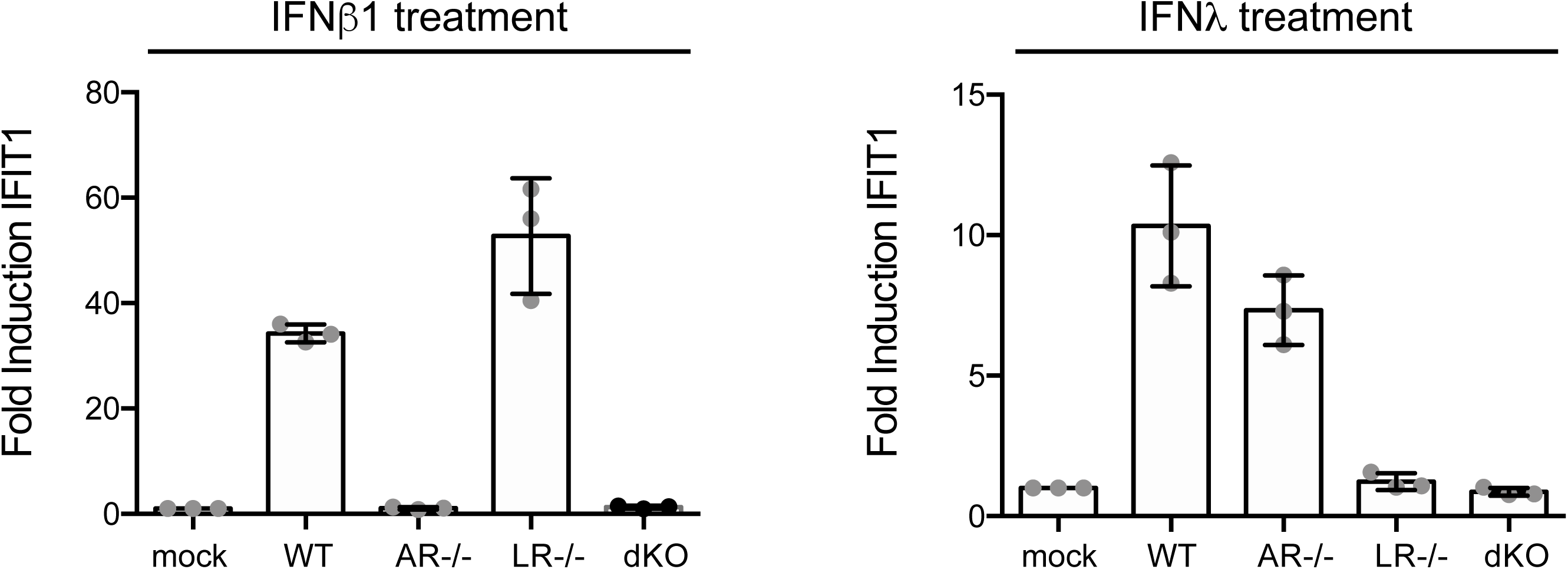
Upregulation of IFIT1 in T84 wild and IFN receptor knock-out cells. Wild type T84 cells and T84 cells depleted of the type I IFN receptor (AR-/-), the type III IFN receptor (LR-/-) or both (dKO) were treated with IFNβ1 and IFNλ. 12 hrs post-treatment, RNA was harvested and the upregulation of IFIT1 was evaluated by q-RT-PCR. N=3. Error bars indicate standard deviation.

**Figure S3.**
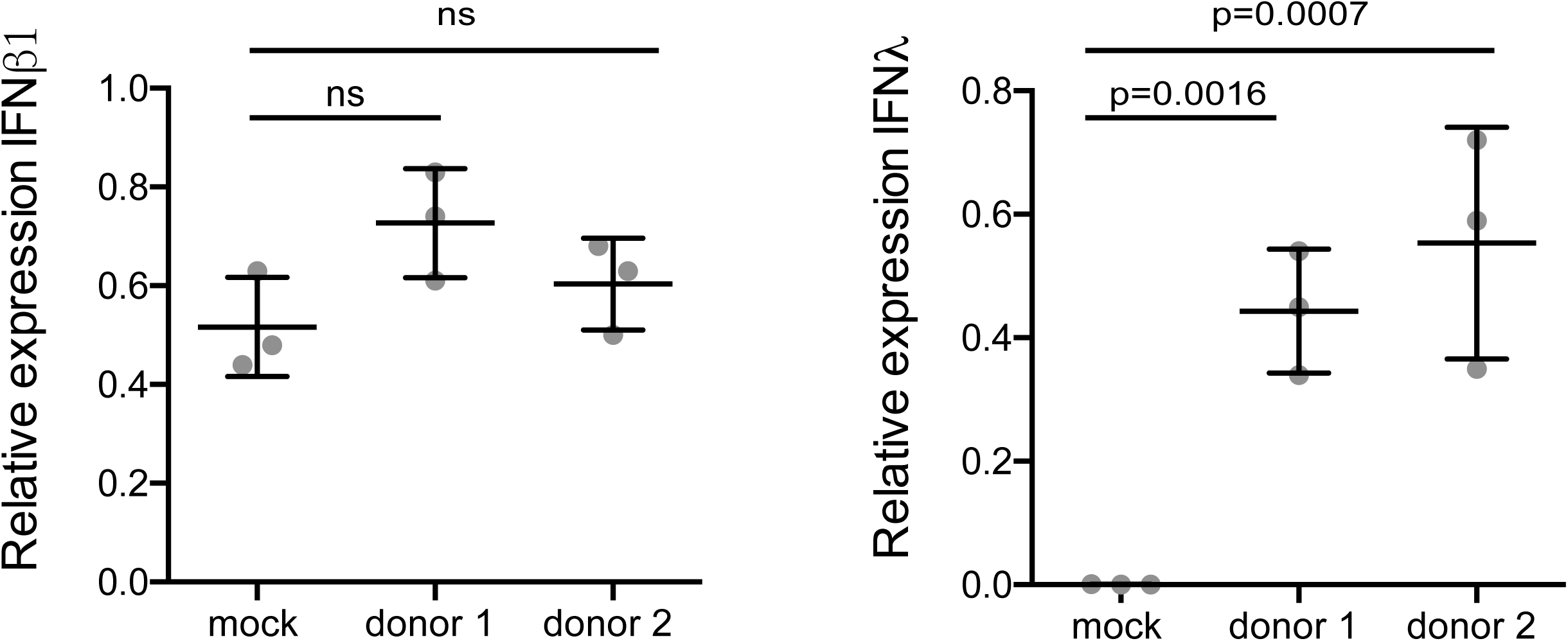
Normalized fold of IFNβ1 and IFNλ induction in colon organoids. Colon organoids were infected with SARS-CoV-2. 24 hpi the upregulation of IFNβ1 and IFNλ was evaluated by q-RT-PCR. N=3. Same data as fig. 4G but values are expressed as normalized to the TBP housekeeping gene. Error bars indicate standard deviation. P values were determined by unpaired t-test.

**Figure S4.**
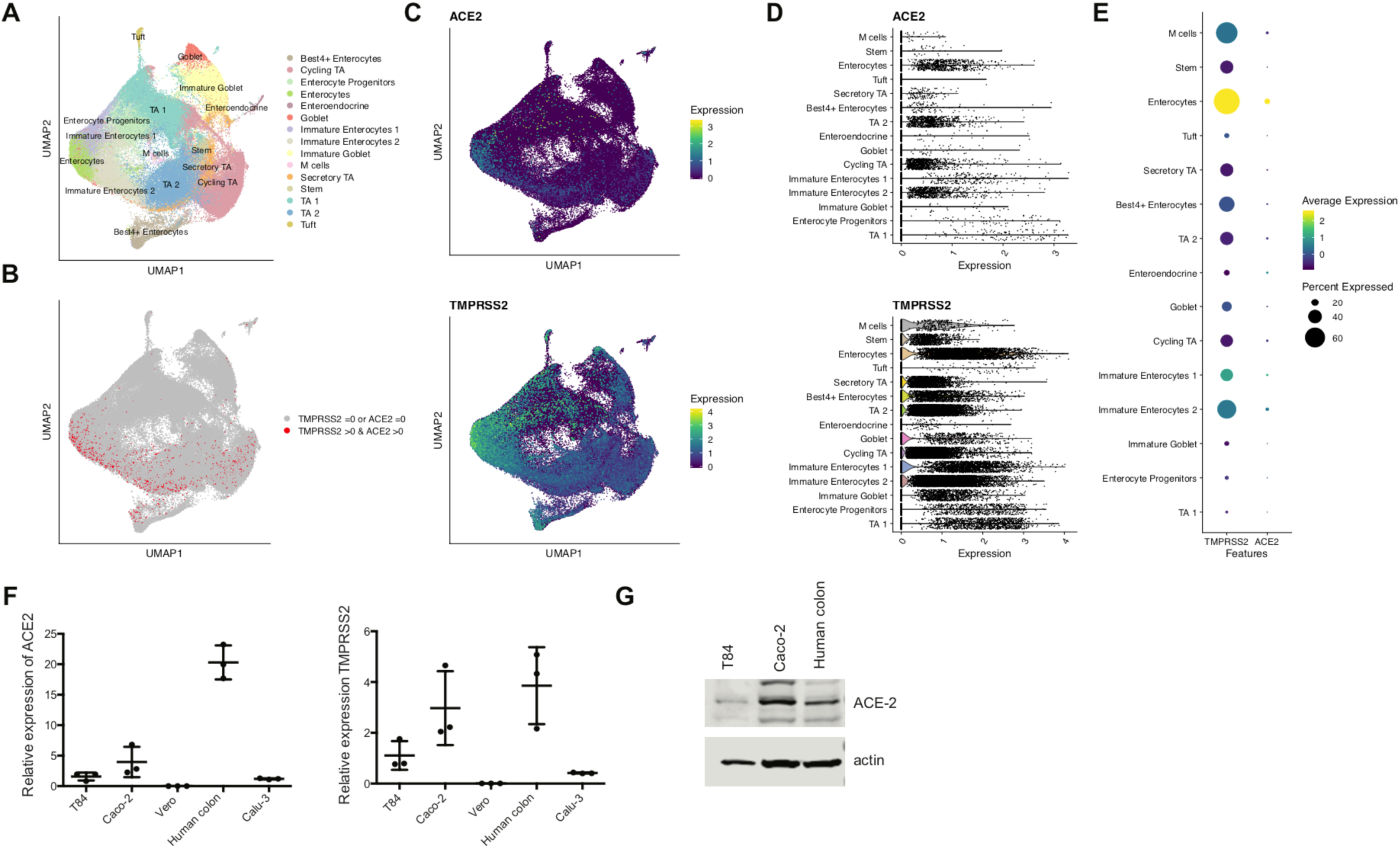
Expression of ACE2 and TMPRSS2 in Human Colon Epithelial cells. (A) Uniform manifold approximation and projection (UMAP) of 99,107 cells from (Smillie et al., 2019) colored by their cell type. (B) UMAP visualization, cells co-expressing ACE2 and TMPRSS2 are highlighted. (C) UMAP visualization, coloured by the normalized expression of ACE2 and TMPRSS2. (D) Expression values of ACE2 and TMPRSS2 in the epithelial cell types of the colon. (E) Dot plot of ACE2 and TMPRSS2 for each cell type. Dot size represents percentage of cells expressing the gene, and colour intensity average expression across the cell type. (F) q-RT-PCR analysis of the levels of ACE-2 and TMPRSS2 in human colon carcinoma cells T84, and Caco-2 cells, Vero E6 cells, human lung adenocarcinoma Calu-3 cells and human colon organoids. Expression is normalized to housekeeping gene TBP. N=3. (G) Western blot of ACE-2 from T84, Caco-2 and human colon organoids. Actin is used as a loading control. Representative image is shown.

## References

Blanco-Melo, D., Nilsson-Payant, B.E., Liu, W.-C., Møller, R., Panis, M., Sachs, D., Albrecht, R.A., and tenOever, B.R. 2020. SARS-CoV-2 launches a unique transcriptional signature from in vitro, ex vivo, and in vivo systems (Microbiology).

Chu, H., Chan, J.F.-W., Wang, Y., Yuen, T.T.-T., Chai, Y., Hou, Y., Shuai, H., Yang, D., Hu, B., Huang, X., et al. Comparative replication and immune activation profiles of SARS-CoV-2 and SARS-CoV in human lungs: an ex vivo study with implications for the pathogenesis of COVID-19. Clin. Infect. Dis.

Crotta, S., Davidson, S., Mahlakoiv, T., Desmet, C.J., Buckwalter, M.R., Albert, M.L., Staeheli, P., and Wack, A. 2013. Type I and Type III Interferons Drive Redundant Amplification Loops to Induce a Transcriptional Signature in Influenza-Infected Airway Epithelia. PLOS Pathog. 9, e1003773.

Cui, J., Li, F., and Shi, Z.-L. 2019. Origin and evolution of pathogenic coronaviruses. Nat. Rev. Microbiol. 17, 181–192.

Ettayebi, K., Crawford, S.E., Murakami, K., Broughman, J.R., Karandikar, U., Tenge, V.R., Neill, F.H., Blutt, S.E., Zeng, X.-L., Qu, L., et al. 2016. Replication of Human Noroviruses in Stem Cell-Derived Human Enteroids. Science 353, 1387–1393.

Harak, C., and Lohmann, V. 2015. Ultrastructure of the replication sites of positive-strand RNA viruses. Virology 479–480, 418–433.

Harcourt, J., Tamin, A., Lu, X., Kamili, S., Sakthivel, S.Kumar., Murray, J., Queen, K., Tao, Y., Paden, C.R., Zhang, J., et al. 2020. Isolation and characterization of SARS-CoV-2 from the first US COVID-19 patient (Microbiology).

Hoffmann, M., Kleine-Weber, H., Schroeder, S., Krüger, N., Herrler, T., Erichsen, S., Schiergens, T.S., Herrler, G., Wu, N.-H., Nitsche, A., et al. 2020. SARS-CoV-2 Cell Entry Depends on ACE2 and TMPRSS2 and Is Blocked by a Clinically Proven Protease Inhibitor. Cell 0.

Leung, W.K., To, K., Chan, P.K.S., Chan, H.L.Y., Wu, A.K.L., Lee, N., Yuen, K.Y., and Sung, J.J.Y. 2003. Enteric involvement of severe acute respiratory syndrome-associated coronavirus infection1. Gastroenterology 125, 1011–1017.

Lokugamage, K.G., Hage, A., Schindewolf, C., Rajsbaum, R., and Menachery, V.D. 2020. SARS-CoV-2 sensitive to type I interferon pretreatment. BioRxiv 2020.03.07.982264.

Lu, H., Stratton, C.W., and Tang, Y.-W. 2020. Outbreak of pneumonia of unknown etiology in Wuhan, China: The mystery and the miracle. J. Med. Virol. 92, 401–402.

Lukassen, S., Lorenz Chua, R., Trefzer, T., Kahn, N.C., Schneider, M.A., Muley, T., Winter, H., Meister, M., Veith, C., Boots, A.W., et al. 2020. SARS-CoV-2 receptor ACE2 and TMPRSS2 are primarily expressed in bronchial transient secretory cells. EMBO J. n/a.

Mantlo, E., Bukreyeva, N., Maruyama, J., Paessler, S., and Huang, C. 2020. Potent Antiviral Activities of Type I Interferons to SARS-CoV-2 Infection. BioRxiv 2020.04.02.022764.

Moriyama, M., Hugentobler, W.J., and Iwasaki, A. 2020. Seasonality of Respiratory Viral Infections. Annu. Rev. Virol. 7, null.

Mossel, E.C., Huang, C., Narayanan, K., Makino, S., Tesh, R.B., and Peters, C.J. 2005. Exogenous ACE2 Expression Allows Refractory Cell Lines To Support Severe Acute Respiratory Syndrome Coronavirus Replication. J. Virol. 79, 3846–3850.

Paules, C.I., Marston, H.D., and Fauci, A.S. 2020. Coronavirus Infections—More Than Just the Common Cold. JAMA 323, 707–708.

Pervolaraki, K., Stanifer, M.L., Münchau, S., Renn, L.A., Albrecht, D., Kurzhals, S., Senís, E., Grimm, D., Schröder-Braunstein, J., Rabin, R.L., et al. 2017. Type I and Type III Interferons Display Different Dependency on Mitogen-Activated Protein Kinases to Mount an Antiviral State in the Human Gut. Front. Immunol. 8, 459.

Pervolaraki, K., Rastgou Talemi, S., Albrecht, D., Bormann, F., Bamford, C., Mendoza, J.L., Garcia, K.C., McLauchlan, J., Höfer, T., Stanifer, M.L., et al. 2018. Differential induction of interferon stimulated genes between type I and type III interferons is independent of interferon receptor abundance. PLoS Pathog. 14, e1007420.

Pott, J., Mahlakõiv, T., Mordstein, M., Duerr, C.U., Michiels, T., Stockinger, S., Staeheli, P., and Hornef, M.W. 2011. IFN-λ determines the intestinal epithelial antiviral host defense. Proc. Natl. Acad. Sci. 108, 7944–7949.

Qi, F., Qian, S., Zhang, S., and Zhang, Z. 2020. Single cell RNA sequencing of 13 human tissues identify cell types and receptors of human coronaviruses. Biochem. Biophys. Res. Commun.

Smillie, C.S., Biton, M., Ordovas-Montanes, J., Sullivan, K.M., Burgin, G., Graham, D.B., Herbst, R.H., Rogel, N., Slyper, M., Waldman, J., et al. 2019. Intra- and Inter-cellular Rewiring of the Human Colon during Ulcerative Colitis. Cell 178, 714-730.e22.

Stanifer, M.L., Rippert, A., Kazakov, A., Willemsen, J., Bucher, D., Bender, S., Bartenschlager, R., Binder, M., and Boulant, S. 2016. Reovirus intermediate subviral particles constitute a strategy to infect intestinal epithelial cells by exploiting TGF-β dependent pro-survival signaling. Cell. Microbiol. 18, 1831–1845.

Stanifer, M.L., Mukenhirn, M., Muenchau, S., Pervolaraki, K., Kanaya, T., Albrecht, D., Odendall, C., Hielscher, T., Haucke, V., Kagan, J.C., et al. 2020. Asymmetric distribution of TLR3 leads to a polarized immune response in human intestinal epithelial cells. Nat. Microbiol. 5, 181–191.

Targett-Adams, P., Boulant, S., and McLauchlan, J. 2008. Visualization of Double-Stranded RNA in Cells Supporting Hepatitis C Virus RNA Replication. J. Virol. 82, 2182–2195.

Venkatakrishnan, A.J., Puranik, A., Anand, A., Zemmour, D., Yao, X., Wu, X., Chilaka, R., Murakowski, D.K., Standish, K., Raghunathan, B., et al. 2020. Knowledge synthesis from 100 million biomedical documents augments the deep expression profiling of coronavirus receptors. BioRxiv 2020.03.24.005702.

Wang, L., and Zhang, Y. 2016. Animal Coronaviruses: A Brief Introduction. In Animal Coronaviruses, L. Wang, ed. (New York, NY: Springer), pp. 3–11.

Wang, Q., Vlasova, A.N., Kenney, S.P., and Saif, L.J. 2019. Emerging and re-emerging coronaviruses in pigs. Curr. Opin. Virol. 34, 39–49.

Wang, W., Xu, Y., Gao, R., Lu, R., Han, K., Wu, G., and Tan, W. 2020. Detection of SARS-CoV-2 in Different Types of Clinical Specimens. JAMA.

Wölfel, R., Corman, V.M., Guggemos, W., Seilmaier, M., Zange, S., Müller, M.A., Niemeyer, D., Jones, T.C., Vollmar, P., Rothe, C., et al. 2020. Virological assessment of hospitalized patients with COVID-2019. Nature.

Wong, S.H., Lui, R.N., and Sung, J.J. Covid-19 and the Digestive System. J. Gastroenterol. Hepatol. n/a.

Wu, C., Zheng, S., Chen, Y., and Zheng, M. (2020a). Single-cell RNA expression profiling of ACE2, the putative receptor of Wuhan 2019-nCoV, in the nasal tissue (Infectious Diseases (except HIV/AIDS)).

Wu, Y., Guo, C., Tang, L., Hong, Z., Zhou, J., Dong, X., Yin, H., Xiao, Q., Tang, Y., Qu, X., et al. (2020b). Prolonged presence of SARS-CoV-2 viral RNA in faecal samples. Lancet Gastroenterol. Hepatol. 0.

Xiao, F., Tang, M., Zheng, X., Liu, Y., Li, X., and Shan, H. 2020. Evidence for gastrointestinal infection of SARS-CoV-2. Gastroenterology S0016508520302821.

Xing, Y.-H., Ni, W., Wu, Q., Li, W.-J., Li, G.-J., Wang, W.-D., Tong, J.-N., Song, X.-F., Wing-Kin Wong, G., and Xing, Q.-S. 2020. Prolonged Viral Shedding in Feces of Pediatric Patients with Coronavirus Disease 2019. J. Microbiol. Immunol. Infect. S1684118220300815.

Xu, H., Zhong, L., Deng, J., Peng, J., Dan, H., Zeng, X., Li, T., and Chen, Q. (2020a). High expression of ACE2 receptor of 2019-nCoV on the epithelial cells of oral mucosa. Int. J. Oral Sci. 12, 1–5.

Xu, Y., Li, X., Zhu, B., Liang, H., Fang, C., Gong, Y., Guo, Q., Sun, X., Zhao, D., Shen, J., et al. (2020b). Characteristics of pediatric SARS-CoV-2 infection and potential evidence for persistent fecal viral shedding. Nat. Med. 1–4.

Ye, L., Schnepf, D., Becker, J., Ebert, K., Tanriver, Y., Bernasconi, V., Gad, H.H., Hartmann, R., Lycke, N., and Staeheli, P. 2019. Interferon-λ enhances adaptive mucosal immunity by boosting release of thymic stromal lymphopoietin. Nat. Immunol. 20, 593–601.

Zhao, Y., Zhao, Z., Wang, Y., Zhou, Y., Ma, Y., and Zuo, W. 2020. Single-cell RNA expression profiling of ACE2, the receptor of SARS-CoV-2 (Bioinformatics).

Zhou, J., Li, C., Zhao, G., Chu, H., Wang, D., Yan, H.H.-N., Poon, V.K.-M., Wen, L., Wong, B.H.-Y., Zhao, X., et al. 2017. Human intestinal tract serves as an alternative infection route for Middle East respiratory syndrome coronavirus. Sci. Adv. 3, eaao4966.

